# Tissue Oxygenation Dynamics During Seizure to Spreading Depolarization in Rat Brain

**DOI:** 10.1101/2025.09.17.676836

**Authors:** Jiayang Liu, Bruce J. Gluckman

## Abstract

**Objective:** Spreading depolarization (SD) is a phenomenon underlying various neurological conditions, including epilepsy. Researchers have suspected that local tissue oxygenation breakdown induces spontaneous SD. In this study, we investigated the relationship between spontaneous epileptic seizures and SD, with a focus on the role of local tissue oxygenation during the transition from seizure to seizure-associated SD.

**Methods:** We applied a long pulse voltametric method to characterize local tissue oxygenation and extracellular space volume in the hippocampus (HC) of freely moving epileptic rats (male 6, female 3). Recordings were performed during the normal state of vigilance, spontaneous seizures, and seizure-associated SD events, as well as during their transitions.

**Results:** No significant breakdown in local tissue oxygenation of HC was detected before SD onset during the seizure to SD transition. In contrast, a decreased ECS volume in the HC was observed before SD onset during this transition.

**Significance:** Using a novel electrochemical approach in freely behaving rats with intact cerebral autoregulation, we demonstrate that ECS shrinkage, rather than breakdown of local tissue oxygenation, plays a leading role in SD initiation during seizure to SD transition. These findings refine our understanding of the mechanisms driving seizure-associated SD and suggest that ECS dynamics may represent an important therapeutic target in epilepsy and other SD-associated neurological disorders

## Introduction

The human brain, although it occupies only about 2% of the body’s mass, accounts for about 20% of the body’s oxygen consumption for resting metabolism (1), and Na^+^-K^+^-ATPase and other ATP-dependent pumps consume a large portion of that 20% to maintain homeostasis of the ion gradients across cell membranes. Arterial blood flow transports oxygen to brain tissues. As blood flows through capillaries, oxygen molecules are continuously released from the red blood cell (RBC) hemoglobin, diffuse across the RBC membrane into the blood plasma, cross the capillary wall, and passively diffuse in the extracellular space (ECS) to the cells in the nearby tissue. The ECS is the narrow microenvironment that surrounds every cell of the central nervous system and occupies about 20% of brain tissue (2). It is usually described and measured by the ECS volume fraction *α*, and the ECS tortuosity *λ* (3).

For neurons in the ECS to integrate signal processing and maintain resting energy metabolism, a continuous and sufficient availability of oxygen, along with proper regulation of the brain tissue oxygenation, is critical. The term spreading depolarization (SD) describes the breakdown of this homeostasis (4). SD describes spectrum of waves in the central nervous system and is characterized by an almost complete breakdown of ion gradients (5), neuronal depolarization block (6), loss of electrical activity (7), and cell swelling and distortion of dendritic spines (8). In the ECS, SD is recorded as a high-amplitude negative DC shift of the slow potential (9). During SD, neurons cannot fire because of sustained depolarization blocks causing an almost complete electrical silence named spreading depression in the cortex of epileptic rabbits (7). In other words, spreading depression is a consequence or epiphenomenon of SD due to a depolarization block of neuronal activity (4).

SD has been reported as the phenomenon underlying migraine aura (10–12) and appears sufficient to activate trigeminal nociception and sustain it for durations consistent with the migraine attack (13). SD is also closely associated with subarachnoid hemorrhage, intracranial hemorrhage (10, 14, 15), stroke, traumatic brain injury (16, 17), epilepsy and epileptogenesis (15, 18), and sudden unexpected death in epilepsy (19).

Experimentally, SD can be induced directly by depolarizing the neurons via sodium and/or calcium channel activation, e.g., administration of glutamate or potassium, or indirectly by depolarizing the neurons through Na/K-ATPase activity reduction, such as ischemia, hypoxia, and hypoglycemia. Electrical or mechanical stimulation has also been adopted to induce SD (20). Considering the significance of SD, a wide range of techniques, including gradient-echo magnetic resonance imaging (MRI), diffusion-weighted MRI, and optical intrinsic signal (OIS) imaging, have been implemented to study SD-associated effects like cell swelling, cerebral blood volume/flow (CBV/CBF), and hemoglobin concentration fluctuation. KCl-evoked SD in rats produced propagating MR signal increases due to venous oxygenation increase (21). SD-induced cell swelling reflected in tissue water apparent diffusion coefficients decreases and negatively correlates with blood oxygenation level (BOLD) signal (22, 23). The ECS volume shrinkage due to SD-induced cell swelling has also been demonstrated in (24). OIS, which measures reflectance changes in light scattering caused by CBV/CBF, hemoglobin, and cytochrome redox fluctuations during ion gradient disturbance and cell swelling, has been used to study SD in both normal and pathological conditions (25, 26).

SD consumes a large amount of oxygen, dramatically interferes with brain tissue oxygenation level, and represents a high-energy challenge for brain tissue (27). The tissue oxygenation dynamics have been studied in different SD models (8, 28–31). Recent slice-level studies using two-photon ATP sensors demonstrate a substantial, often transient drop in neuronal ATP during SD, underscoring risks in ischemic contexts (32).

A key limitation of previous studies is their reliance on induced SD models, which are often conducted under artificial conditions such as ischemia or hypoxia. These models do not allow for the assessment of SD initiation or propagation in the presence of intact neurovascular regulation. Additionally, some studies draw conclusions about causation from correlation; for example, they may assume that hypoxia-induced SD accurately reflects normal seizure pathophysiology, a potential interpretative error highlighted in (33). Theoretical modeling work (34) suggests that tissue oxygenation plays an important role in understanding the interactions between spontaneous seizures and SD. The tissue oxygenation dynamics in seizures and SD events have been studied on animal brain slices (35). However, these works lack physiological autoregulation, intact metabolism, and neurovascular coupling regulation.

Finally, the ECS itself is increasingly recognized as a dynamic regulator of brain health. Recent studies show the ECS not as a passive conduit but a structurally heterogeneous, actively regulated space modulating molecular diffusion and neuronal function, making ECS modulation a potential therapeutic frontier for neurological disorders (36).

Using continuous long-term recordings in the tetanus toxin (TeTX) rat model of temporal lobe epilepsy (TLE), we previously demonstrated synchronized recordings of electrophysiology and electrochemistry during different state transitions, including seizure to SD transitions (37). In this study, we investigate the **causal mechanisms** underlying these transitions with intact blood flow autoregulation and metabolic control, with particular emphasis on the initiation of SD. As illustrated in **Figure 1**, in the TeTX model, when an animal enters a spontaneous seizure from a normal state of vigilance (SOV), one of two trajectories follows: seizure termination or progression into SD. The mechanisms determining this divergence remain unclear. We examine tissue oxygenation dynamics across normal SOV, peri-seizure, and peri-SD transitions. By applying long pulse voltammetry (LPV), we decouple the effects of bulk oxygen concentration from the effective diffusion coefficient, enabling indirect characterization of ECS volume fraction changes during these transitions.

**Figure 1:**
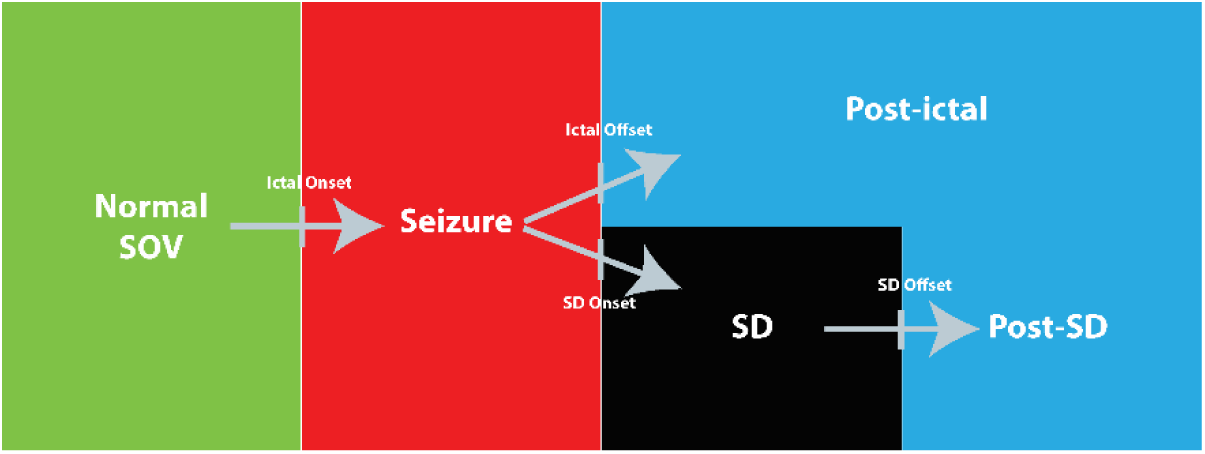
State transition pathways in the tetanus toxin rat model of temporal lobe epilepsy. From a normal state of vigilance (SOV, green), a rat enters a spontaneous seizure (red), which can lead to one of two trajectories: the rat can go into a postictal state (blue) or go into spreading depolarization (SD, black) followed by a post-SD state (blue).

## Methods

### One dimension model of oxygen diffusion and consumption in brain tissue

We used the tetanus toxin (TeTX) model of temporal lobe epilepsy (TLE), in which presynaptic inhibitory neurotransmitter release is disrupted, leading to spontaneous seizures and seizure-associated SD (38). This model provides a suitable platform to study SD initiation and seizure–SD interactions under intact autoregulation. To monitor tissue oxygenation, we performed long-term, continuous biopotential and electrochemical recordings using constant potential amperometry (CPA) and long pulse voltammetry (LPV). CPA with microelectrodes yields a mixed oxygenation signal reflecting both bulk oxygen concentration (*C_B_*) and the effective diffusion coefficient (*D_eff_*), but it assumes *D_eff_* is constant and does not allow the two to be separated (39). In contrast, the LPV enables decoupling of this mixed signal: the initial current peak reflects local oxygen concentration, whereas the steady-state flat tail corresponds to the CPA-like mixed measure. From these signals, relative changes in *D_eff_* can be extracted, providing an indirect estimate of ECS volume dynamics.

Oxygen delivery to brain tissue depends on cardiorespiratory function and hemoglobin binding (∼98% of total oxygen) (Pittman, 2011). Previous studies on diffusion models (40–42) assume uniform oxygen consumption, constant diffusion, and even-spaced capillaries. However, these assumptions break down during seizures and SD, when (1) *C_B_* fluctuates with CBF/CBV and hemoglobin oxygenation, and (2) *D_eff_* is altered by changes in ECS volume fraction (*α*) and tortuosity (*λ*). To account for these dynamics, we implemented a one-dimensional (1D) diffusion model of brain tissue oxygenation as shown in **Figure 2A**. Arteries deliver oxygen bound to hemoglobin within red blood cells (RBCs). As blood flows into capillaries, oxygen is released from hemoglobin, diffuses into the interstitial fluid, and then from the bulk tissue (near capillaries) to the oxygen-sensing site at the electrochemical working electrode (WE) tip.

**Figure 2:**
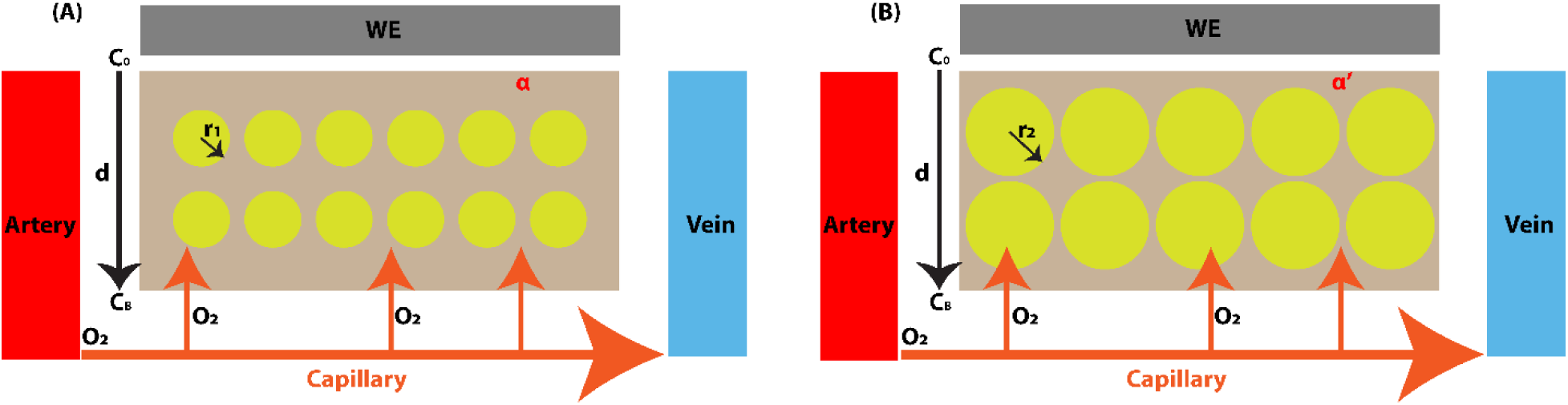
One-dimensional model of oxygen diffusion in brain tissue. Oxygen molecules (O₂) carried by red blood cells (RBCs) enter capillaries from the artery (red) to vein (blue), are released from hemoglobin, and continuously diffuse into the interstitial fluid. (**A**) normal SOV, (**B**) pathological states. *WE*: Pt working electrode. *r_1_/r_2_*: cell (yellow) radius.*C_0_*: local oxygen concentration at the WE surface. *C_B_*: bulk oxygen concentration near the capillary. *d*: Nernst diffusion thickness. *α/α’*: ECS volume fraction.

Because the WE tip area is much larger than individual cells, axial oxygen diffusion can be neglected. We assume a uniform oxygen consumption rate across tissue. The average distance between the WE surface (*x = 0*) and the nearest capillary is 25*µ*m on average in vivo, and typically 100 *µ*m from the bath when recording in vitro (34). This distance (*d*) is assumed to remain constant during each animal’s recordings, regardless of behavioral state, which included rapid eye movement (REM) sleep, non-REM (NREM) sleep, wake, seizure, and SD event. Within brain tissue, cell centers are assumed fixed; only cell radius (*r*) increases or decreases, leading to changes in ECS volume fraction (*α*), while tortuosity (*λ*) is assumed to be constant. Consequently, the *D_eff_* is modeled as dependent on *α*.

Equations when we consider oxygen reduction only at the electrode to tissue interface: Current from Fick’s Law (Mass Transport). The oxygen reduction Faradaic current at the electrode surface (or diffusion current) is governed by the diffusion and for 1D model we have:

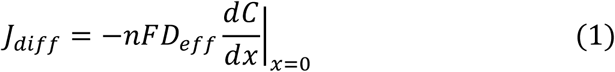

Where: *D_eff_* is oxygen diffusion coefficient, *n* is number of electrons transferred, *F* is Faraday constant, *d* is Nernst diffusion layer thickness, *C_B_* is bulk oxygen concentration at *x = d*.

### Voltametric methods solutions

We applied two voltametric methods to the model: constant potential amperometry (CPA) and long pulse voltammetry (LPV). In CPA, a constant bias potential (-0.65V), i.e., a big overpotential, is applied and its current response:

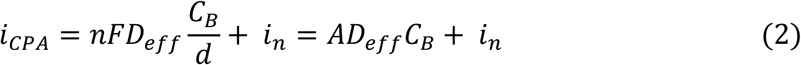

Where: *i_n_* represents non-Faradaic current and other measurement noise, *A* is a geometric factor. From equation (2), we know that the CPA measure when overpotential is big, is a mixed result depending on the bulk tissue oxygen concentration *C_B_*, and the effective oxygen diffusion coefficient *D_eff_*. The CPA provides a way to learn the tissue oxygenation dynamics determined by *C_B_* and *D_eff_*. However, the separation of the mixed result is difficult, especially during pathological states (e.g., seizures or SD events) have not been studied. In our model, we used LPV to decouple *C_B_* and *D_eff_* and characterize them separately.

The bias potential waveform used in LPV is shown in **Figure 3**. The potential *E_1_* lies in the region where an equilibrium state is held. *E_2_* is set in the “mass-transfer-limited” region which makes the local oxygen concentration on the working electrode surface go nearly to zero rapidly due to high oxygen reduction kinetics. As the bias potential changes from *E_1_* to *E_2_*, a large overpotential is applied and the initial current peak value (*pv*) is dominantly dependent on the initial oxygen concentration before overpotential is applied which is the bulk oxygen concentration *C_B_*, so we can get:

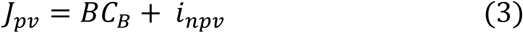

**Figure 3:**
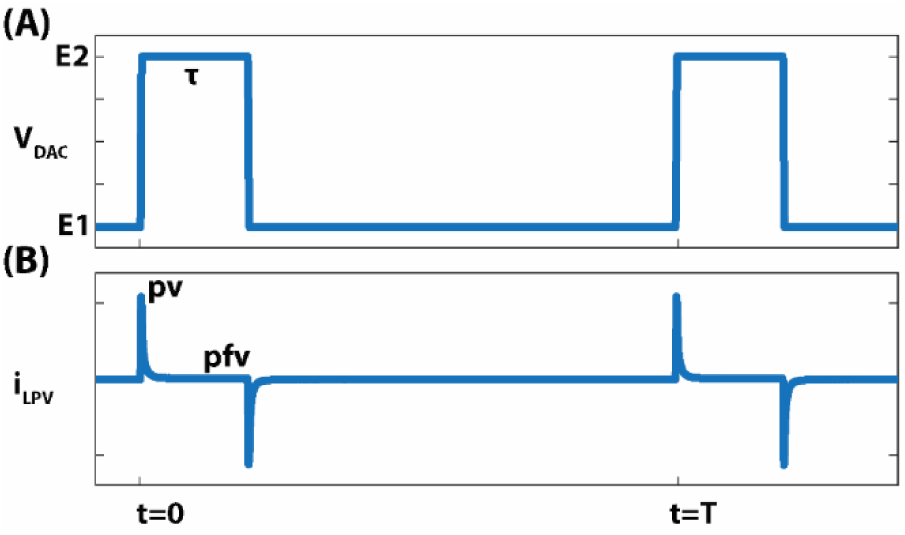
The long pulse voltammetry (LPV) waveform and current response example. (**A**) Bias potential waveform. (**B**) response current, *pv*: LPV current response’s peak value, *pfv*: LPV current response’s peak flat value. T: period. τ:pulse duration.

Where: *i_npv_* represents non-Faradaic current and other measurement noise, *B* is a geometric factor.

After the initial current peak, current flows continue maintaining the fully oxygen reduced condition at potential *E_2_* on the working electrode surface which becomes the CPA process as discussed above, and we get the current peak flat value (*pfv*):

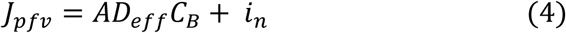

To extract *D_eff_*, we make the following assumptions:

1. Within each state (e.g., NREM, seizure, or SD), *D_eff_* = *D̄ + ΔD, C_B_ = C̄_B_ + ΔC_B_* and *ΔC_B_* and *ΔD_eff_* are dependent on its state and independent of each other.
2. *i_n_* and *i_npv_* are dependent on its state and independent of each other.

Based on equation (3) and (4), through the ratio between the covariance of *J_pv_* and *J_pfv_* and the variance of *J_pv_*.

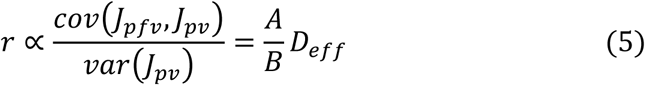

From equation (5), we can use the ratio value *r* to characterize the *D_eff_*. For the same animal, *r* is state-dependent. From one state to another state (e.g., NREM to REM, or Normal SOV to seizure), we can extract *D_eff_* by doing ensemble averages of the ratios with respect to transition times. <u>Alternately, for peri-transition time ensembles, we take the variance and covariance at particular offset times with respect to the transition time directly across the events ensemble</u>. The latter method has the advantage that the sample sizes are larger and the variances in concentration are in practice larger than the local noise terms.

### Relationship between the extracellular space volume fraction and the effective oxygen diffusion coefficient

One property of cell membranes is that they are barriers to the permeation of macro-molecules. However, studies of micro-molecules, e.g., oxygen molecules, have shown that the cell membrane is not a complete barrier for oxygen transportation (43–49). These studies focus on the process of oxygen being transported across the cell membrane from the ECS into the cell and the oxygen concentration difference across the cell membrane. In (39), these studies have been reviewed and explained in detail. However, there are two oxygen diffusion pathways in the tissue: one is through the ECS with a diffusion coefficient *D_1_*, and the other through the cells with a diffusion coefficient *D_2_*, as shown in **Figure 2A**. The two pathways have different effective oxygen diffusion coefficients, and their relative contributions are determined by cell morphology under different brain states. When the animal’s state changes, e.g., from seizure to SD, we assume the cellular center remains fixed while morphological changes are reflected by cell radius changes (**Figure 2B**). Because this morphological change is primarily driven by water flux, we assume that *D_1_* and *D_2_* remain constant. Assuming uniform oxygen solubility in brain tissue, the effective oxygen diffusion coefficient (*D_eff_*) can be expressed as:

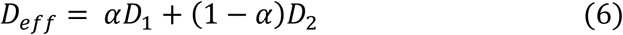

Where *α* is the ECS volume fraction and occupies ∼20% of the total volume (2). From equation (6), changes in *D_eff_* can be used to indirectly characterize alterations in ECS volume fraction *α*.

### Animal surgery and care

All experimental protocols were approved by the Institutional Animal Care and Use Committee. We used Long-Evans rats (6 males and 3 females) from Charles River Laboratories, LLC, or ENVIGO, weighing 250 to 500 grams.

For depth local field potential (LFP) recordings in the hippocampus, we employed 50 µm diameter micro-reaction chamber (µRC) electrode (50) pairs with tips 125 ∼ 250 µm apart dorsally. Stainless steel screws were employed for measuring ECoG. For electrochemical recordings using the CPA or LPV, we employed a 200 μm diameter platinum (Pt) wire with PET (polyethylene terephthalate) plastic film coating as the working electrode (WE), a custom-made Ag/AgCl electrode as the reference electrode (RE), and a screw electrode as the counter electrode (CE). The Ag/AgCl electrode was created by inserting a 100 µm diameter silver wire into a 155 µm diameter polyamide tube filled with Ag/AgCl ink (CI-4001 Silver/Silver Chloride/Vinyl, Nagase America LLC).

All surgeries were conducted under deep anesthesia using Ketamine (90 mg/kg) and Xylazine (15 mg/kg). A heating pad was utilized under the animal to maintain a constant body temperature of 37°C. Rats were secured in a stereotaxic frame with ear bars and given pre-operative Buprenorphine Ethiqa XR (3.25 mg/kg) for pain relief. Lidocaine (<5 mg/kg) was injected subcutaneously at the incision site. Burr holes were drilled with an electric drill based on stereotaxic coordinates relative to bregma (Mouse Atlas, Paxinos and Franklin, 2001 (Paxinos 2001)) as shown in **Figure 4**. Specific targets and naming convention included: hippocampus at HAR (hippocampal anterior right), HPL (hippocampal posterior left), HPR (hippocampal posterior right) to provide measurements of the hippocampal local field potentials (LFP). Stainless steel screws measuring ECoG are at EFL (ECoG frontal left), EFR (ECoG frontal right), EAL (ECoG anterior left), EAR (ECoG anterior right), EPL (ECoG posterior left), and EPR (ECoG posterior right). O_2_-WE (oxygen-sensing working electrode), O_2_-Ref (oxygen-sensing reference electrode).

**Figure 4:**
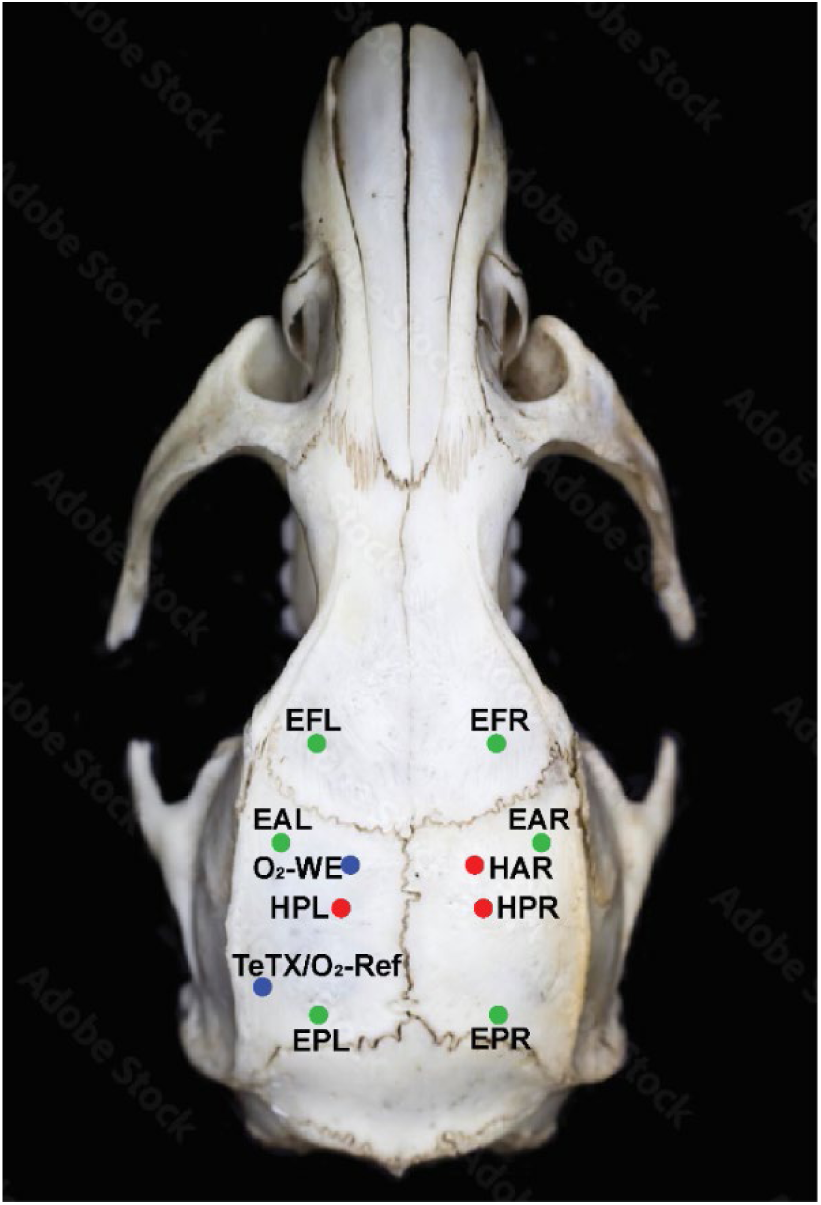
Stereotaxic coordinate of electrodes. All coordinates were bregma-referenced (AP, ML, DV). Red dots show coordinates of depth electrodes for local field potential (LFP) measurements in the hippocampal. HAR (hippocampal anterior right): -2.5mm, +2.0mm, -3.2mm. HPL/R (hippocampal posterior left/right): -3.9mm, ±2.2mm, -2.88mm. Blue dots show oxygen sensing related electrodes. O_2_-WE (Oxygen-sensing working electrode): -2.5mm, - 2.0mm, -3.2mm. TeTX/O_2_-Ref (TeTX injection site/Oxygen sensing reference): -5.51mm, - 5.35mm, -6.1mm. Green dots show coordinates of stainless-steel screw electrodes measuring ECoG. EFL/R (EcoG frontal left/right): 2mm, ±3mm. EAL/R (EcoG anterior left/right): 1.5mm, ±4mm. EPL/R (EcoG posterior left/right): -6.5mm, ±4mm. Oxygen sensing reference electrode is implanted after toxin injection at the same site to minimize additional cortical damage. Skull figure adopted from (96).

Rats were prepared under the TeTX model (51) and has been used to induce seizures in mice and rats (52). Researchers have characterized the mechanisms of action of the toxin as well as seizure development and progression (51, 53–56). Usually, after the toxin injection, the animal will begin to experience spontaneous seizures within 10 days. The procedures for toxin injection and electrode implantation are outlined in (57). Briefly, 10 to 13 nano-grams of tetanus toxin (Santa Cruz Biotechnology, CAS 676570-37-9) dissolved in 1.3 microliters phosphate-buffered saline (PBS) mixed with 2% bovine serum albumin (BSA) are injected into the rat’s left ventral hippocampus (AP -5.15, ML -5.35, DV -6.1 mm) through a 30-gauge flexible cannula over 15 minutes with additional 30 minutes for tissue relaxation. The WE for oxygen sensing was implanted to target the hippocampus anterior left (HAL), and the custom-made Ag/AgCl RE was placed in the cortex along the same trajectory that was used for toxin injection, minimizing further damage to the cortex. LFP recording depth electrodes, ECoG screw electrodes, WE, and RE were secured in place on the skull and electrically isolated via dental cement. After surgery, rats were returned to individual standard autoclave-ready cages with free access to food and water and maintained at a 12-hour light-dark cycle with lights on between 7 am and 7 pm. We allowed a seven-day post-surgery recovery before initiating recordings.

### Oxygen-sensing working electrode calibration

The *in vitro* oxygen-sensing calibration setup for the working electrode (WE) is described in (37). In short, we implemented a three-electrode electrochemical cell with a platinum (Pt) wire WE, an Ag/AgCl pellet RE, and a Pt plate CE. Oxygen content was known by first preparing two separate PBS solutions from the same sample, one was saturated with air, and the other with virtually no oxygen, formed by bubbling Nitrogen (N_2_) through it. Three electrodes were submerged in the air-saturated solution, followed by incremental addition of the N₂-saturated solution. The experiment was conducted at room temperature with the PBS solution’s pH value at 7.48 (pH Meter, Model P771. Anaheim Scientific). The CPA and LPV waveforms were provided by the electrochemical instrument described in (37). The resulting oxygen concentration is derived from standard stoichiometric calculations.

The CPA calibration result is shown in **Figure 5A**. In brain tissue, the oxygen concentration, the partial pressure of oxygen (*pO_2_*), and the oxygen tension are mutually related values that, in principle, can be derived from one another. The oxygen concentration is expressed in moles per liter (mol/l) and is used for measuring dissolved oxygen. The *pO_2_* reflects the amount of free oxygen molecules and equals the pressure that oxygen would exert if it occupied the space by itself (39). The oxygen concentration is expressed in moles per liter (mol/l) in this study for measuring dissolved oxygen. The CPA calibration is well-approximated by a linear fit over the physiological range from ∼0.09 mM/L (the tissue oxygen concentrations) to ∼0.23 mM/L (the arterial oxygen concentration) (58) marked by the purple and blue lines. The LPV peak flat value (*pfv*), which is comparable to the CPA measurement as discussed in the modeling section, exhibits a similar calibration result (**Figure 5B**). Given that the calibration process occurs in a PBS solution, we assume that the effective oxygen diffusion coefficient remains constant. This assumption has been supported by the peak value (*pv*) calibration results illustrated in **Figure 5C**.

**Figure 5:**
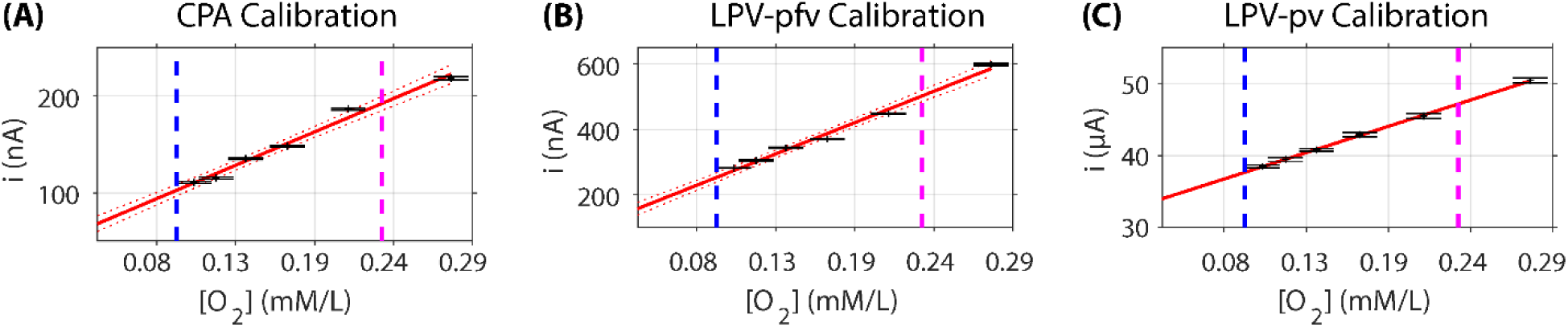
CPA and LPV calibration of oxygen-sensing electrodes. Linear fitting results of current response versus oxygen concentration (X-axis, oxygen concentration; Y-axis, current). (**A**) CPA calibration result. (**B**) LPV peak flat value (*pfv*) calibration result. (**C**) LPV peak value (*pv*) calibration result. Each data point was derived from the addition of N2-saturated PBS solution. For each addition, we recorded data for approximately 5 minutes, and each value was calculated from the middle 80 seconds of this recording to eliminate any transients associated with the addition process.

### Data analysis and statistics

Recorded data include LFP from *µ*RC electrodes targeting the hippocampus, ECoG from screw electrodes over the cortex, three-axis head acceleration signal, and tissue oxygenation signals using the CPA and LPV. Data is processed offline using custom-written MATLAB (MathWorks Inc.) programs for re-referencing, filtering, spectral analysis, and behavior annotation. Hippocampal LFPs are band-pass filtered at 0.5 ∼ 125Hz to highlight field potentials and seizure dynamics. Seizure and SD detections have been described in (37). Seizures were detected by a stereotypical increase in spectral power that initiates with a sentinel spike followed by a burst of 9 ∼ 16 Hz hippocampal spikes (59), spreads through the cortex, and ends with a sharp decrease in spectral power. Seizures that are shorter than 10 seconds and spaced apart less than 10 minutes are excluded. **SD detections** were done from depth electrode measures referenced to a cortical screw electrode. SD events were detected from at least one depth electrode (i.e., from at least one out of HAR, HPL, or HPR) signals low pass filtered with a cutoff at 0.5 Hz. SD onset was defined by a downward crossing of a 7.5 mV threshold in the signal with respect to the value 3 seconds before. SD offset was then detected by an upward crossing of a threshold defined as 2 mV above the SD onset potential. **The propagation speed of SD** was determined by dividing the physical distance between electrodes HAR and HPR by the difference in time indices corresponding to the downward SD crossing in these two channels. Only animals with ‘clean’ signals on HAR and HPR were used for SD propagation speed calculation. Normal SOV was marked as one of the three states: REM sleep characterized by a spectral peak in the theta (4 ∼ 7 Hz) frequency band of hippocampal LFP and by an absence of acceleration except during brief muscle twitches; NREM sleep characterized by maximal power in the delta (0.5 ∼ 4 Hz) frequency band and by an absence of acceleration; wake characterized by the accelerometer activity (60, 61). As demonstrated in **Figure 1**, transitions include NREM to REM, NREM to wake, normal SOV to seizure to normal SOV, normal SOV to seizure to SD.

For each animal, CPA response currents were preprocessed by removing recordings with big artifacts or filling ‘nan’ for small artifacts. After preprocessing, for each transition, after a low-pass filter at 10 Hz, the CPA response time course was baseline normalized, and down-sampled from 1 kHz to 20 Hz. The baseline normalization was done by subtracting the 60 s pre-state mean and then dividing the 60 s pre-state mean to manifest the dynamics change and make the time-course trace start from 0%. The normalized CPA response time course median was displayed with quantile from a 60 s pre-state to a post-state, with a display cutoff at 60 s for the SD events, and 30 s for other post-states. For statistical analysis, the CPA response changes within pre-state and post-state were averaged (without the cutoff) across all transitions. Compared to CPA, LPV current responses are “discrete” pulses. After the same preprocessing as CPA, we extracted two values from each pulse as shown in **Figure 3**. The current pulse peak value (*pv*) comes from the bias potential transition that turns on the oxygen reduction reaction and depends on the bulk oxygen concentration (*C_B_*). The current peak flat value (*pfv*) sampled ∼10ms before the next transition, which is comparable with the CPA current, depends on the bulk tissue oxygenation (*C_B_*), and the effective oxygen diffusion coefficient (*D_eff_*). Using *pv* and *pfv*, we extracted a ratio value (*rv*) that characterizes the change of *D_eff_* as detailed in the model session. All LPV response features (i.e., *pv, pfv, rv*) from all transitions were baseline z-score normalized to get a time course from a 60 s pre-state to post-state transition, with a display cutoff at 60s for the SD period, and 30 s for other post-states. For statistical analysis, the two-sample Kolmogorov-Smirnov test (kstest2 function in Matlab) was used to detect significant changes. 5 s pre-state data and 5 s post-state data were chosen with an offset 5 s or 10 s to transition onset time, e.g., -10 s ∼ -5 s before transition onset time at 0, and 5 s ∼ 10 s after onset time. All data are shown as mean ± standard deviation, and significance level was set at p < 0.001 or p < 0.05.

## Results

We aim to examine long-term local tissue oxygenation dynamics in vivo in freely moving animals across normal SOV transitions, peri-ictal transitions, and peri-SD transitions. Specifically, we want to determine whether local tissue oxygenation plays a leading or determining role in transitions from normal SOV to seizure, and from seizure to SD, or whether it primarily reflects a reactive response. Additionally, we aim to investigate whether there is evidence of a significant breakdown in local oxygen regulation during these transitions. Furthermore, we want to decouple and characterize the dynamics of the effective oxygen diffusion coefficient (*D_eff_*) during these events, alongside changes in ECS volume fraction. In previous study, we demonstrated CPA measures combined with electrophysiological recordings (37). In this study, we applied a new LPV measure. A 200-second representative recording episode from hippocampal depth electrodes, ECoG screws, and the LPV measures from oxygen-sensing working electrode are shown in **Figure 6**. Band-pass filtered ECoG and hippocampal (HC) LFP measurements are shown in **Figure 6A**. Low pass filtered HC LFP measurements demonstrate the initiation and propagation of SD in the HC, which are not detectable in the band-pass filtered data (**Figure 6B**). The SD propagation speed is: 4.3 ± 2 mm/min (mean ± std, 44 SD events), consistent with previous studies on SD propagation speed (9, 62, 63). The LPV measure, using a 1 Hz waveform with a 200 ms pulse duration, provides a means to decouple local oxygen concentration and the *D_eff_* through *pv* and *pfv* measures, as detailed in the modeling section. LPV *pv* (red dots, left Y-axis) and LPV *pfv* (blue dots, right Y-axis) are shown in **Figure 6C**.

**Figure 6:**
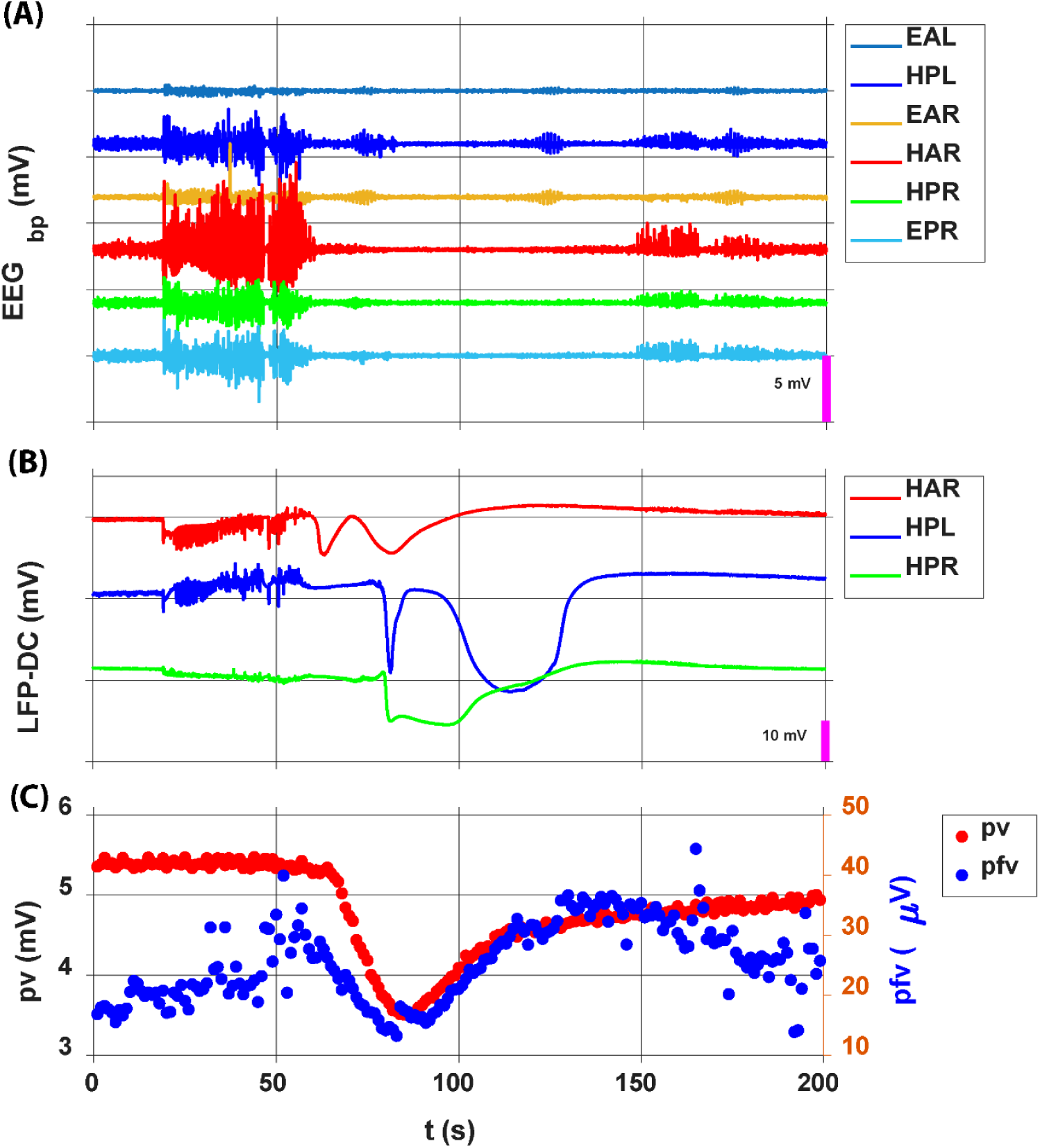
Representative recordings of hippocampal LFP, ECoG, and LPV during a seizure to SD transition. A 200-second episode showing simultaneous recordings from hippocampal depth electrodes, ECoG screws, and the oxygen-sensing working electrode. **(A)** Band-filtered ECoG and hippocampal LFP measurements. **(B)** Low pass filtered hippocampal LFP measurements. **(C)** LPV current response from the oxygen-sensing working electrode. LPV *pv* (red dots, left Y-axis), LPV *pfv* (blue dots, right Y-axis).

### CPA current and LPV feature dynamics during normal state of vigilance transitions

Using CPA in rats, previous studies have demonstrated that brain tissue oxygen levels increase during wakefulness and REM sleep and decline during NREM sleep (64, 65). Consistent with this, another study showed that the transition from NREM to wake is accompanied by a marked increase in oxygen, reflecting enhanced inspired oxygen and ventilation associated with wakefulness (66). Similarly, previous work has reported that arousing stimuli elicit rapid elevations in brain oxygen, further supporting the sensitivity of CPA to state- and stimulus-dependent changes in oxygen availability (67).

Shown in **Figure 7** are the dynamics of CPA current and LPV features during normal SOV transitions, from NREM sleep to wake, and from NREM sleep to REM sleep. During the NREM to wake transition, CPA current shows a significant increase (**Figure 7A, 7E**). LPV *pfv*, comparable to CPA current, shows a similar significant increase (**Figure 7B, 7F**). LPV *pv*, reflecting local tissue oxygenation, decreases slightly (< 1%) without significance (**Figure 7C, 7G**). LPV *rv*, reflecting the effective diffusion coefficient (*D_eff_*), remains stable (**Figure 7D**), indicating no ECS volume fraction change.

**Figure 7:**
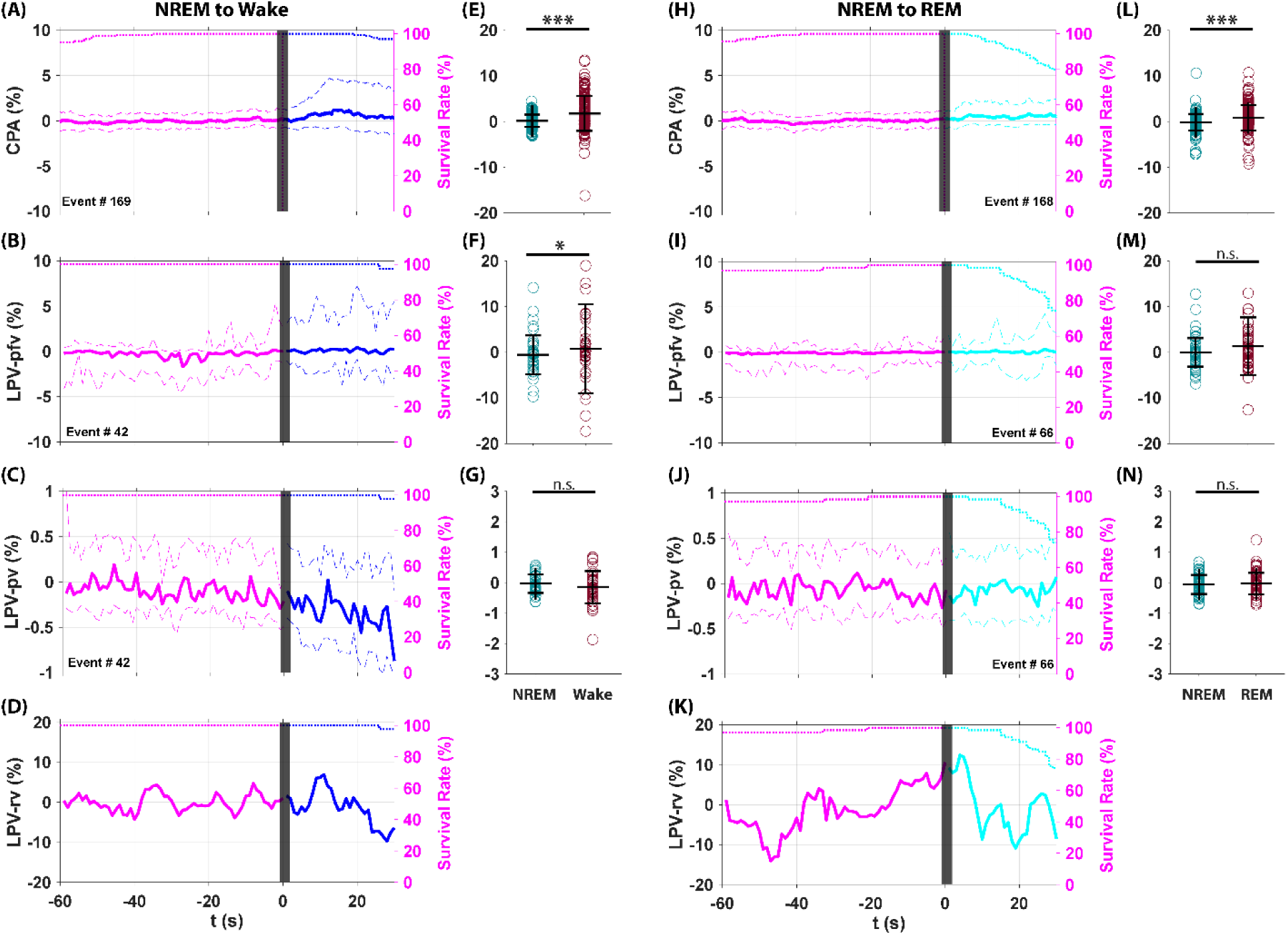
CPA current and LPV feature dynamics during normal state of vigilance transitions. Normal state of vigilance (SOV) transitions include: NREM sleep to REM sleep transition, and NREM sleep to wakefulness transition. REM onset and wake onset are at t = 0 (black bar). Each trace shows the median ± quantiles across observed transitions (left Y-axis, solid line ± dashed line, Magenta: NREM, Blue: Wake, Cyan: REM). State survival curves (right Y-axis, dotted line, Magenta: NREM, Blue: Wake, Cyan: REM) indicate the probability of remaining in a given state, helping the interpretation of ensemble-averaged dynamics. During the NREM to wake transition, CPA current showed a significant increase (**A, E**). LPV *pfv*, comparable to CPA current, showed a similar significant increase (**B, F**). LPV *pv*, reflecting local tissue oxygenation, decreased slightly (<1%) without significance (**C, G**). LPV *rv*, reflecting the effective diffusion coefficient (*D_eff_*), remained stable (**D**). During the NREM to REM transition, CPA current again showed a significant increase (**H, L**). LPV *pfv* showed a similar but non-significant trend (**I, M**). LPV *pv* remained unchanged (**J, N**), and LPV *rv* also remained stable (**K**). Statistical analyses confirmed significant CPA current changes (**L**), whereas LPV *pfv*, though showed similar dynamic trend, didn’t show significant change, possibly due to limited sample size (**M**).

During the NREM to REM transition, CPA current again shows a significant increase (**Figure 7H, 7L**). LPV *pfv* shows a similar but non-significant trend (**Figure 7I, 7M**). LPV *pv* remains unchanged (**Figure 7J, 7N**), and LPV *rv* also remains stable (**Figure 7K**), indicating no ECS volume fraction change. During NREM to REM transition, statistical analyses confirm a significant CPA current increase (**Figure 7L**), whereas LPV *pfv,* though shows similar dynamic trend, does not show significant change, possibly due to limited sample size (**Figure 7M**).

In summary, consistent with previous reports, we observed significant increases in oxygen levels during normal SOV transitions using CPA. However, LPV *pfv* measurements show no significant changes in local oxygenation during normal SOV transitions. LPV *rv*, reflecting *D_eff_*, remained unchanged, indirectly indicating no significant alterations in ECS volume fraction during normal SOV transitions.

### CPA current and LPV feature dynamics during transitions from normal SOV to seizure and from seizure (without SD associated) back to normal SOV

Previous studies have shown that seizures profoundly alter local oxygenation dynamics. Optical and oxygen-sensitive probe recordings consistently report a transient “initial dip” in oxygen availability within the ictal focus, reflecting a rapid surge in metabolic demand during seizure that briefly exceeds supply, whereas recordings from surrounding areas show increased oxygenation without this dip (68–70). These findings highlight the spatiotemporal heterogeneity of seizure-related vascular responses and suggest that peri-ictal regions undergo distinct oxygenation dynamics.

The CPA and LPV recordings are shown in Figure 8. During the transition from normal SOV to seizure and back to normal SOV, CPA current first increased significantly (**Figure 8A, 8E**) and then decreased significantly after seizure offset during recovery (**Figure 8H, 8L**). LPV *pfv*, which is comparable to CPA current, showed a similar trend with a significant increase (**Figure 8B, 8F**) followed by a non-significant decrease, likely due to limited sample size and higher noise levels (**Figure 8I, 8M**). LPV *pv*, reflecting local tissue oxygenation, remained unchanged during seizure (**Figure 8C, 8G**) but showed a significant decrease after seizure offset (**Figure 8J, 8N**). LPV *rv*, reflecting the *D_eff_*, decreased during seizure and increased after seizure offset to wakefulness (**Figure 8D, 8K**).

**Figure 8:**
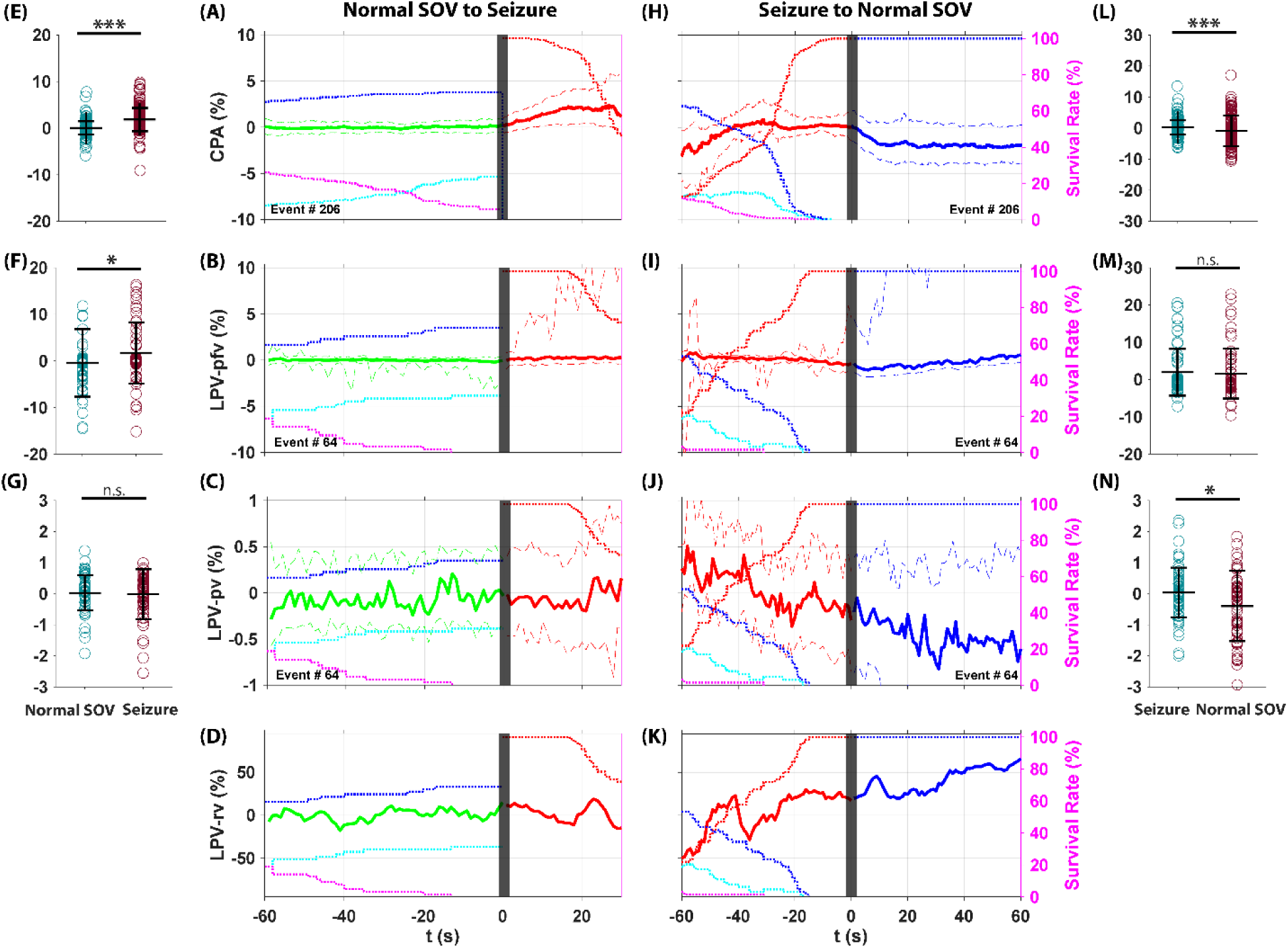
CPA current and LPV feature dynamics during transitions from normal SOV to seizure and from seizure back to normal SOV. Each trace shows the median ± quantiles across observed transitions (left Y-axis, solid line ± dashed line, Green: normal SOV. Red: seizure. Blue: wake). State survival curves (right Y-axis, dotted line, Red: seizure, Magenta: NREM, Blue: Wake, Cyan: REM) indicate the probability of remaining in a given state, helping the interpretation of ensemble-averaged dynamics. Seizure onset and seizure offset are at t = 0 (black bar). CPA current first increased significantly (**A, E**) and then decreased significantly after seizure offset during recovery (**H, L**). LPV *pfv*, which is comparable to CPA current, showed a similar trend with a significant increase (**B, F**) followed by a non-significant decrease, likely due to limited sample size and higher noise levels (**I, M**). LPV *pv*, reflecting local tissue oxygenation, remained unchanged during seizure (**C, G**) but showed a significant decrease after seizure offset (**J, N**). LPV *rv*, reflecting the *D_eff_*, decreased during seizure and increased after seizure offset to wakefulness (**D, K**).

Our oxygen-sensing WE was implanted in the area close but not at the TeTX injection site (i.e., ictal focus). We demonstrate an overall increased ictal oxygenation without the initial dip in either CPA or LPV recording, consistent with previous findings in seizure foci surrounding areas (68–70). Importantly, LPV *pv* measurements showed no evidence of local oxygenation breakdown before seizure onset, indicating that peri-ictal oxygenation followed the transition from normal SOV to seizure without an anticipatory deficit. After seizure offset, we detected a postictal decrease in local oxygenation, consistent with earlier studies reporting prolonged hypoxia (71). In addition, LPV *rv* measurements indirectly showed an ictal ECS shrinkage (cell swelling), which aligns with prior studies: seizures in mice have been shown to reduce ECS volume by ∼ 35% (72), and a ∼ 15% shrinkage has been reported during spontaneous epileptiform discharges in trauma-injured rat neocortex (73).

### CPA current and LPV features dynamics during transitions from normal SOV to seizure and to seizure-associated SD

As illustrated in **Figure 1**, there are two pathways in our TeTX model following the seizure: the rat can either return to a normal SOV or enter a SD. CPA current and LPV features during the transitions from normal SOV to seizure, and from seizure to seizure-associated SD, are shown in **Figure 9**.

**Figure 9:**
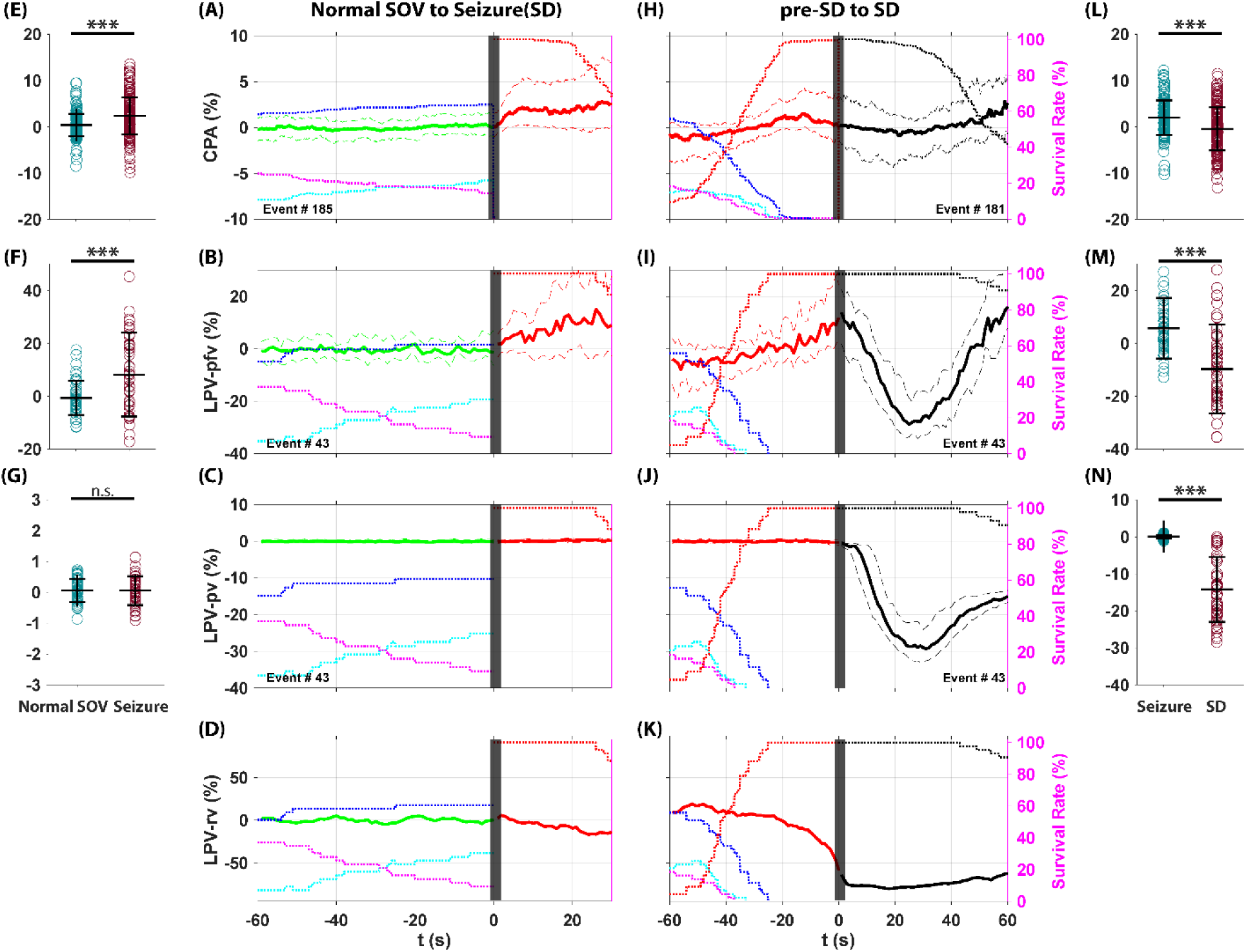
CPA current and LPV features dynamics during transitions from normal SOV to seizure and to seizure-associated SD. Each trace shows the median ± quantiles across observed transitions (solid line ± dashed line, Green: normal SOV. Red: seizure. Black: SD). State survival curves (right Y-axis, dotted line, Black: SD, Red: seizure, Magenta: NREM, Blue: Wake, Cyan: REM) indicate the probability of remaining in a given state, helping the interpretation of ensemble-averaged dynamics. Seizure onset and SD onset (seizure offset) are at t = 0 (black bar). The CPA current showed a significant increase during the transition from normal SOV to seizure (**A, E**) and a significant decrease during SD, recovering as the rate of SD survival decreased (**H, L**). The LPV *pfv* also showed similar dynamics across this transition: it first increased from normal SOV to seizure (**B, F**) and then decreased significantly during SD, returning to baseline as SD survival rates declined (**I, M**). However, the LPV *pv* did not show significant changes during the transition from normal SOV to seizure (**C, G**), but after the onset of SD, the *pv* decreased significantly (**J, N**). The LPV *rv* began to decline as seizure survival increased before SD onset (**D**) and continued to drop by as much as 80% during SD (**K**).

The CPA current showed a significant increase during the transition from normal SOV to seizure (**Figure 9A, 9E**) and a significant decrease during SD, recovering as the rate of SD survival decreased (**Figure 9H, 9L**). The LPV *pfv*, which is comparable to CPA current, also showed similar dynamics across this transition: it first increased from normal SOV to seizure (**Figure 9B, 9F**) and then decreased significantly during SD, returning to baseline as SD survival rates declined (**Figure 9I, 9M**). However, the LPV *pv* did not show significant changes during the transition from normal SOV to seizure (**Figure 9C, 9G**), but after the onset of SD, the *pv* decreased significantly (**Figure 9J, 9N**). This indicates that local oxygenation does not break down and behaves as a following role during normal SOV to seizure transition. Decreased local oxygenation during SD aligns with previous studies, which show that direct tissue recordings of partial oxygen pressure (*pO₂*) confirm that SD can lead to a rapid drop in local oxygen levels, occasionally reaching near-anoxic conditions, before partially recovering depending on vascular reactivity and the baseline perfusion state (8, 19).

During normal SOV to seizure transition, the LPV *rv* began to decline as seizure survival increased, importantly, this decline occurred before the onset of SD (**Figure 9D**). After the onset of SD, the LPV *rv* continued to drop by as much as 80% during SD (**Figure 9K**). This result demonstrates that the ECS shrinkage plays a leading role in the transition from seizure to SD. This result after SD onset is also consistent with previous study describing a neuronal swelling during SD events (8) which shrinks the ECS volume.

In summary, during the peri-SD transition, CPA and LPV *pfv* showed similar patterns, with a significant increase followed by a significant decrease. However, CPA averages differed by only a few percent (**Figure 9A, 9H**), whereas LPV *pfv* showed larger changes of 15–25% (**Figure 9B, 9I**). We demonstrated that ECS volume dropped by as much as 80% during SD (**Figure 9K**), indicating that *D_eff_* cannot be assumed constant. Given that LPV *pv* has a much higher signal-to-noise ratio and is therefore the more reliable measure, the large ECS volume change implies that both CPA and LPV *pfv* were corrupted by ECS volume shrinkage, which explains the difference between the two measurements.

### The extracellular space dominates the oxygen diffusion in the tissue

The ECS volume change affects the *D_eff_* as detailed in the model session. From one state to another state, e.g., from seizure to SD, the *α* changes to *α’*. The center of the cell remains unchanged, and the morphological change is shown with a larger *r* as shown in **Figure 2B**. We get:

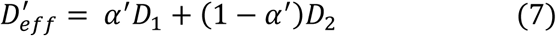

Where *α’* is the ECS volume fraction in the new state. Oxygen diffusion pathways distance from local to bulk, *d*, is assumed to be constant. We set *α* = 0.2 during normal SOV (2), and make an extreme assumption that *α’ = 0* during SD events (**Figure 9K**). From equation (6) and (7), we get *D_2_ = 21 × D_1_*, which demonstrates that the <u>ECS is the dominate oxygen diffusion pathway</u> in the tissue and is altered dramatically by the ECS volume fraction change during SD events.

## Discussion

The level of tissue oxygenation reflects the dynamic balance between oxygen delivery, utilization, tissue reactivity, and morphology under physiological conditions (66). Constant potential amperometry (CPA) has been used for in vivo oxygen sensing. However, CPA-based studies often assume that the effective diffusion coefficient (*D_eff_*) is constant, are unable to decouple the effects of local oxygen concentration and the changes of *D_eff_* (39). We introduce a one-dimensional model of oxygen diffusion in brain tissue under normal SOV, seizures, and seizure-associated spreading depolarization (SD), as well as their transitions. The CPA is current response is a mixed signal with both local oxygen concentration (*C₀*) and *D_eff_*. Using a long pulse voltammetry (LPV) we can decouple these components, with the LPV peak flat value (*pfv*) corresponding to a measure comparable to CPA current, and the LPV peak value (*pv*) reflecting *C₀*. Using *pfv* and *pv*, we extract a ratio value (*rv*), characterizing changes in the *D_eff_* directly and ECS volume fraction indirectly during different state transitions, importantly, during spontaneous seizure to seizure-associated SD event transition.

From CPA and LPV *pfv* measures, we demonstrate increased recordings during transitions from NREM to both wake and REM states (**Figure 7A, 7B, 7H, 7I**), consistent with prior works showing elevated oxygen during wakefulness and REM relative to NREM (64, 66). These increases reflect enhanced ventilation and oxygen intake upon arousal, together with local neurovascular adjustments that support increased neuronal activities (67, 74, 75). Interestingly, despite these CPA-detected changes, LPV *pv* and *rv* measurements demonstrated no significant change in local tissue oxygenation or in ECS volume during normal SOV transitions (**Figure 7C, 7D, 7J, 7K**). This indicates that temporary increases in oxygen during behavioral arousal in normal SOV are significantly smaller or “buffered” by a baseline homeostatic mechanism with considerable metabolic demands, which helps maintain stable local oxygenation and ECS volume under normal physiological conditions (74, 75). The baseline metabolism buffering is also consistent with the tight coupling between local metabolism and cerebral blood flow, which stabilizes oxygen delivery across states (76, 77). Previous studies have indirectly demonstrated that although rapid stimulation-induced oxygen dynamics can be detected with high temporal resolution methods such as CPA or fast oxygen electrodes, baseline oxygenation remains stable across physiological conditions (65, 78).

Our findings during peri-ictal and peri-SD highlight dynamic changes in local oxygenation and ECS volume. During seizures, CPA current and LPV *pfv* demonstrated significant increases, consistent with previous studies (**Figure 8A, 8B, 8H, 8I**). Notably, LPV *pv* showed no significant breakdown of local oxygenation before seizure onset and during ictal period, indicating that peri-ictal elevated metabolic demand was compensated by increased cerebral oxygen delivery through intact neurovascular coupling and local oxygenation was preserved despite the surge in neuronal activity (68–70). In parallel, we demonstrated decreased ECS volume (or cell swelling) during seizures (**Figure 8D, 8K)**, consistent with prior studies of activity-dependent ECS constriction. Such shrinkage likely facilitates the accumulation of ECS potassium and neurotransmitters, amplifying excitability and promoting seizure propagation (79–83).

During the transition from seizures to SD, both CPA current and LPV *pfv* showed significant decreases (**Figure 9A, 9B, 9H, 9I**). However, LPV *pv*, reflecting local oxygenation, showed different dynamics: remained stable before SD onset (**Figure 9C**) then showed a significant decrease during SD (**Figure 9J**). This finding demonstrates two key points: first, local oxygenation does not break down during seizures before SD onset, suggesting that impaired oxygen supply is not the primary trigger for SD initiation. Second, the collapse of local oxygenation during SD reflects a near-anoxic state, consistent with direct tissue *pO₂*recordings that show dramatic oxygen depletion during SD (8, 19) and with the elevated metabolic demand required for recovery (84, 85). This severe oxygen decline also reflects a failure of neurovascular coupling during SD (86). It has been demonstrated indirectly in stroke patients using intraoperative hybrid photoacoustic and ultrasonic imaging to monitor blood oxygenation during SD (87). Similar results have been reported in multiple rodent models. Previous rat study showed that SD was accompanied by an initial hypoperfusion (∼ 75% of baseline), followed by transient hyperemia (∼ 220% of baseline) and subsequent oligemia (28). Researchers reported a large drop in tissue *pO₂* after SD onset with a less pronounced decrease in CBF, followed by significant increases in both tissue *pO₂* and CBF (30, 31). In a mouse study, researchers showed that worsening mismatches between oxygen supply and consumption drive the initiation and propagation of SD (88). The high metabolic demand during SD arises from massive, near-complete depolarization of neurons and glia, collapse of ionic gradients, and recruitment of ATP-dependent pumps to restore homeostasis (4, 89). When combined with impaired or spatially restricted vascular responses, this overwhelming demand-supply mismatch results in sustained hypoxia.

Researchers described neuronal swelling during SD events (8). The ECS volume has been reported to decrease from 20 % to approximately 5 % because of the water influx due to ion changes, leading to intracellular hyperosmolality (5, 24, 90, 91). As detailed in the model session, the ECS volume faction change alters *D_eff_*, allowing indirect estimation of ECS volume dynamics through *D_eff_* measurements. As shown in **Figure 9D, 9K**, we demonstrate that the ECS volume decrease starts during seizure, before the SD onset, and continues to decrease dramatically during SD up to 80 %, suggesting that significant ECS shrinkage (or cell swelling) may play a leading role in inducing SD during seizures. The restricted ECS diffusion may exacerbate metabolic stress by further limiting oxygen and substrate availability and trigger SD (72, 73).

Together, these results underscore the fundamentally different local oxygenation and ECS volume dynamics of seizures versus SD. Seizures, in intact tissue, appear metabolically sustainable: neurovascular coupling remains preserved, and compensatory hyperemia supports oxygen delivery despite concurrent ECS constriction. In contrast, SD imposes an overwhelming metabolic burden associated with near-complete neuronal and glial depolarization, collapse of ionic gradients, and dramatic ECS shrinkage. This burden overwhelms vascular compensation, leading to profound hypoxia. Our results may help explain why seizures are often reversible, whereas seizure-associated SDs are strongly linked to neuronal injury and adverse clinical outcomes (16, 92).

Our results also demonstrate the ECS is the dominant pathway for oxygen diffusion in brain tissue, consistent with prior theoretical and experimental studies. Classic models of tissue oxygenation first described oxygen diffusion from capillaries into the surrounding extracellular fluid as the primary route for substrate delivery (40). Following studies have shown that ECS volume fraction and tortuosity critically shape the efficiency of oxygen transport (81). Experimental work further supports this view, with measurements of cortical oxygen dynamics indicating that ECS diffusion governs both the spatial extent and temporal resolution of oxygen delivery to active neuronal populations (93). While oxygen can also cross cell membranes and diffuse through the cytoplasm to reach mitochondria (94, 95), this intracellular pathway plays a secondary role compared to the ECS pathway. Our results, showing that ECS shrinkage during seizures and seizure-associated SD strongly constrains oxygen availability, reinforce the view that ECS diffusion is the principal determinant of local oxygen dynamics.

## Author Contributions

JL and BJG designed and constructed the hardware, designed the research, analyzed the data, and wrote the paper.

## Acknowledgement

We thank the participation of Carlos Curay and Chole Melnick who served in implanting the animals.

## Conflict of Interest

## Funding sources

This work is partially supported under NIH awards R01EB019804 and R01EB014641.

## Notes

### Competing Interest Statement

The authors have declared no competing interest.

## Reference

1. Rolfe DF, Brown GC. Cellular energy utilization and molecular origin of standard metabolic rate in mammals. Physiol Rev. 1997;77(3):731–58.

2. Nicholson C, Hrabetová S. Brain Extracellular Space: The Final Frontier of Neuroscience. Biophysical Journal. 2017;113(10):2133–42.

3. Syková E, Nicholson C. Diffusion in brain extracellular space. Physiological Reviews. 2008;88(4):1277–340.

4. Dreier JP. The role of spreading depression, spreading depolarization and spreading ischemia in neurological disease. Nat Med. 2011;17(4):439–47.

5. Kraig RP, Nicholson C. Extracellular Ionic Variations during Spreading Depression. Neuroscience. 1978;3(11):1045–59.

6. Canals S, Makarova I, Largo C, Ibarz JM, Herreras O. Longitudinal depolarization gradients along the somatodendritic axis of CA1 pyramidal cells: A novel feature of spreading depression. Journal of Neurophysiology. 2005;94(2):943–51.

7. Leao AAP. Spreading depression of activity in the cerebral cortex. Journal of Neurophysiology. 1944;7(6):359–90.

8. Takano T, Tian GF, Peng WG, Lou NH, Lovatt D, Hansen AJ, et al. Cortical spreading depression causes and coincides with tissue hypoxia. Nature Neuroscience. 2007;10(6):754–62.

9. Leao AAP. Further Observations on the Spreading Depression of Activity in the Cerebral Cortex. Journal of Neurophysiology. 1947;10(6):409–14.

10. Gorji A. Spreading depression: a review of the clinical relevance. Brain Res Rev. 2001;38(1-2):33–60.

11. Brennan KC, Charles A. Sleep and Headache. Semin Neurol. 2009;29(4):406–18.

12. Pietrobon D, Moskowitz MA. Chaos and commotion in the wake of cortical spreading depression and spreading depolarizations. Nature Reviews Neuroscience. 2014;15(6):379–93.

13. Brennan KC, Pietrobon D. A Systems Neuroscience Approach to Migraine. Neuron. 2018;97(5):1004–21.

14. Dreier JP, Woitzik J, Fabricius M, Bhatia R, Major S, Drenckhahn C, et al. Delayed ischaemic neurological deficits after subarachnoid haemorrhage are associated with clusters of spreading depolarizations. Brain. 2006;129:3224–37.

15. Dreier JP, Major S, Pannek HW, Woitzik J, Scheel M, Wiesenthal D, et al. Spreading convulsions, spreading depolarization and epileptogenesis in human cerebral cortex. Brain. 2012;135:259–75.

16. Lauritzen M, Dreier JP, Fabricius M, Hartings JA, Graf R, Strong AJ. Clinical relevance of cortical spreading depression in neurological disorders: migraine, malignant stroke, subarachnoid and intracranial hemorrhage, and traumatic brain injury. J Cerebr Blood F Met. 2011;31(1):17–35.

17. Basheer N, Varghese JC, Kuruvilla R, Alappat JP, Mathew J, Basheer N. A Prospective Study on the Incidence and Outcome of Cranial Nerve Injuries in Patients with Traumatic Brain Injuries. Indian J Neurotraum. 2021;18(01):45–50.

18. Eickhoff M, Kovac S, Shahabi P, Ghadiri MK, Dreier JP, Stummer W, et al. Spreading depression triggers ictaform activity in partially disinhibited neuronal tissues. Experimental Neurology. 2014;253:1–15.

19. Aiba I, Noebels JL. Spreading depolarization in the brainstem mediates sudden cardiorespiratory arrest in mouse SUDEP models. Sci Transl Med. 2015;7(282):282ra46.

20. Mathew AA, Panonnummal R. Cortical spreading depression: culprits and mechanisms. Exp Brain Res. 2022;240(3):733–49.

21. Gardnermedwin AR, Vanbruggen N, Williams SR, Ahier RG. Magnetic-Resonance-Imaging of Propagating Waves of Spreading Depression in the Anesthetized Rat. J Cerebr Blood F Met. 1994;14(1):7–11.

22. James MF, Smith MI, Bockhorst KHJ, Hall LD, Houston GC, Papadakis NG, et al. Cortical spreading depression in the gyrencephalic feline brain studied by magnetic resonance imaging. J Physiol-London. 1999;519(2):415–25.

23. Latour LL, Hasegawa Y, Formato JE, Fisher M, Sotak CH. Spreading Waves of Decreased Diffusion-Coefficient after Cortical Stimulation in the Rat-Brain. Magnet Reson Med. 1994;32(2):189–98.

24. Mazel T, Richter F, Vargová L, Syková E. Changes in extracellular space volume and geometry induced by cortical spreading depression in immature and adult rats. Physiol Res. 2002;51:S85–S93.

25. Wang Z, Li PC, Luo WH, Chen SB, Luo QM. Peri-infarct temporal changes in intrinsic optical signal during spreading depression in focal ischemic rat cortex. Neuroscience Letters. 2007;424(2):133–8.

26. Zheng Z, Cao Z, Luo J, Lv J. Characterization of Intrinsic Optical Signal during Spreading Depolarization. Neuropsychiatry. 2018;08.

27. Lindquist BE. Spreading depolarizations pose critical energy challenges in acute brain injury. J Neurochem. 2024;168(5):868–87.

28. Ayata C, Shin HK, Salomone S, Ozdemir-Gursoy Y, Boas DA, Dunn AK, Moskowitz MA. Pronounced hypoperfusion during spreading depression in mouse cortex. J Cerebr Blood F Met. 2004;24(10):1172–82.

29. Yuzawa I, Sakadzic S, Srinivasan VJ, Shin HK, Eikermann-Haerter K, Boas DA, Ayata C. Cortical spreading depression impairs oxygen delivery and metabolism in mice. J Cerebr Blood F Met. 2012;32(2):376–86.

30. Sakadzic S, Yuan S, Dilekoz E, Ruvinskaya S, Vinogradov SA, Ayata C, Boas DA. Simultaneous imaging of cerebral partial pressure of oxygen and blood flow during functional activation and cortical spreading depression. Appl Optics. 2009;48(10):D169–D77.

31. Hobbs CN, Johnson JA, Verber MD, Wightman RM. An implantable multimodal sensor for oxygen, neurotransmitters, and electrophysiology during spreading depolarization in the deep brain. Analyst. 2017;142(16):2912–20.

32. Schoknecht K, Baeza-Lehnert F, Hirrlinger J, Dreier JP, Eilers J. Spreading depolarizations exhaust neuronal ATP in a model of cerebral ischemia. Proc Natl Acad Sci U S A. 2025;122(19):e2415358122.

33. Bernard C. On Fallacies in Neuroscience. Eneuro. 2020;7(6).

34. Wei YN, Ullah G, Schiff SJ. Unification of Neuronal Spikes, Seizures, and Spreading Depression. Journal of Neuroscience. 2014;34(35):11733–43.

35. Ingram J, Zhang CF, Cressman JR, Hazra A, Wei YN, Koo YE, et al. Oxygen and seizure dynamics: I. Experiments. Journal of Neurophysiology. 2014;112(2):205–12.

36. Yong Y, Cai, Y., Lin, J., et al. Advancement in modulation of brain extracellular space and unlocking its potential for intervention of neurological diseases. Med-X. 2024;2, 6 (2024).

37. Liu J, Gluckman BJ. A DC-sensitive video/electrophysiology monitoring unit for long-term continuous study of seizures and seizure-associated spreading depolarization in a rat model. bioRxiv. 2025:2025.02.04.635811.

38. Kramer DR, Fujii T, Ohiorhenuan I, Liu CY. Interplay between Cortical Spreading Depolarization and Seizures. Stereot Funct Neuros. 2017;95(1):1–5.

39. Subczynski WK, Swartz HM. EPR Oximetry in Biological and Model Samples. In: Eaton SR, Eaton GR, Berliner LJ, editors. Biomedical EPR, Part A: Free Radicals, Metals, Medicine, and Physiology. Boston, MA: Springer US; 2005. p. 229–82.

40. Krogh A. The number and distribution of capillaries in muscles with calculations of the oxygen pressure head necessary for supplying the tissue. J Physiol-London. 1919;52(6):409–15.

41. Hill AV. The diffusion of oxygen and lactic acid through tissues. P R Soc Lond B-Conta. 1928;109(728):39–96.

42. Krogh A. The Anatomy and Physiology of Capillaries: Yale University Press; 1922.

43. Nicolson P, Roughton FJW. A Theoretical Study of the Influence of Diffusion and Chemical Reaction Velocity on the Rate of Exchange of Carbon Monoxide and Oxygen between the Red Blood Corpuscle and the Surrounding Fluid. Proc R Soc Ser B-Bio. 1951;138(891):241–64.

44. Holland RAB. Kinetics of Combination of O2 and Co with Human Hemoglobin F in Cells and in Solution. Resp Physiol. 1967;3(3):307–+.

45. Chan HC, Glockner JF, Swartz HM. Oximetry in Cells and Tissues Using a Nitroxide-Liposome System. Biochim Biophys Acta. 1989;1014(2):141–4.

46. Glockner JF, Swartz HM, Pals MA. Oxygen Gradients in Cho Cells - Measurement and Characterization by Electron-Spin Resonance. J Cell Physiol. 1989;140(3):505–11.

47. Hu HP, Sosnovsky G, Swartz HM. Simultaneous Measurements of the Intracellular and Extracellular Oxygen Concentration in Viable Cells. Biochim Biophys Acta. 1992;1112(2):161–6.

48. Glockner JF, Norby SW, Swartz HM. Simultaneous Measurement of Intracellular and Extracellular Oxygen Concentrations Using a Nitroxide-Liposome System. Magnet Reson Med. 1993;29(1):12–8.

49. Swartz HM. Measurements of Intracellular Concentrations of Oxygen - Experimental Results and Conceptual Implications of an Observed Gradient between Intracellular and Extracellular Concentrations of Oxygen. Oxygen Transport to Tissue Xv. 1994;345:799–806.

50. Shanmugasundaram B, Gluckman BJ. Micro-reaction chamber electrodes for neural stimulation and recording. Annu Int Conf IEEE Eng Med Biol Soc. 2011;2011:656–9.

51. Jefferys JGR, Walker MC. Tetanus Toxin Model of Focal Epilepsy. Models of Seizures and Epilepsy. 2006:407–14.

52. Mellanby J, George G, Robinson A, Thompson P. Epileptiform syndrome in rats produced by injecting tetanus toxin into the hippocampus. J Neurol Neurosurg Psychiatry. 1977;40(4):404–14.

53. Jefferys JG, Borck C, Mellanby J. Chronic focal epilepsy induced by intracerebral tetanus toxin. Ital J Neurol Sci. 1995;16(1-2):27–32.

54. Sunderam S, Chernyy N, Mason J, Peixoto N, Weinstein SL, Schiff SJ, Gluckman BJ. Seizure modulation with applied electric fields in chronically implanted animals. Conf Proc IEEE Eng Med Biol Soc. 2006;2006:1612–5.

55. Sunderam S, Chernyy N, Peixoto N, Mason JP, Weinstein SL, Schiff SJ, Gluckman BJ. Seizure entrainment with polarizing low-frequency electric fields in a chronic animal epilepsy model. J Neural Eng. 2009;6(4):046009.

56. Sedigh-Sarvestani M, Thuku GI, Sunderam S, Parkar A, Weinstein SL, Schiff SJ, Gluckman BJ. Rapid eye movement sleep and hippocampal theta oscillations precede seizure onset in the tetanus toxin model of temporal lobe epilepsy. J Neurosci. 2014;34(4):1105–14.

57. Sedigh-Sarvestani M, Thuku GI, Sunderam S, Parkar A, Weinstein SL, Schiff SJ, Gluckman BJ. Rapid Eye Movement Sleep and Hippocampal Theta Oscillations Precede Seizure Onset in the Tetanus Toxin Model of Temporal Lobe Epilepsy. Journal of Neuroscience. 2014;34(4):1105–14.

58. Ortiz-Prado E, Dunn JF, Vasconez J, Castillo D, Viscor G. Partial pressure of oxygen in the human body: a general review. Am J Blood Res. 2019;9(1):1–14.

59. Finnerty GT, Jefferys JGR. 9-16 Hz oscillation precedes secondary generalization of seizures in the rat tetanus toxin model of epilepsy. Journal of Neurophysiology. 2000;83(4):2217–26.

60. Sunderam S, Chernyy N, Peixoto N, Mason JP, Weinstein SL, Schiff SJ, Gluckman BJ. Improved sleep-wake and behavior discrimination using MEMS accelerometers. J Neurosci Meth. 2007;163(2):373–83.

61. Bahari F, Ssentongo P, Schiff SJ, Gluckman BJ. A Brain-Heart Biomarker for Epileptogenesis. Journal of Neuroscience. 2018;38(39):8473–83.

62. Herreras O, Somjen GG. Propagation of spreading depression among dendrites and somata of the same cell population. Brain Res. 1993;610(2):276–82.

63. Cozzolino O, Marchese M, Trovato F, Pracucci E, Ratto GM, Buzzi MG, et al. Understanding Spreading Depression from Headache to Sudden Unexpected Death. Front Neurol. 2018;9:19.

64. Dash MB, Tononi G, Cirelli C. Extracellular levels of lactate, but not oxygen, reflect sleep homeostasis in the rat cerebral cortex. Sleep. 2012;35(7):909–19.

65. McHugh SB, Fillenz M, Lowry JP, Rawlins JN, Bannerman DM. Brain tissue oxygen amperometry in behaving rats demonstrates functional dissociation of dorsal and ventral hippocampus during spatial processing and anxiety. Eur J Neurosci. 2011;33(2):322–37.

66. Ortiz-Prado E, Natah S, Srinivasan S, Dunn JF. A method for measuring brain partial pressure of oxygen in unanesthetized unrestrained subjects: the effect of acute and chronic hypoxia on brain tissue PO(2). J Neurosci Methods. 2010;193(2):217–25.

67. Solis E, Jr., Cameron-Burr KT, Kiyatkin EA. Rapid Physiological Fluctuations in Nucleus Accumbens Oxygen Levels Induced by Arousing Stimuli: Relationships with Changes in Brain Glucose and Metabolic Neural Activation. Front Integr Neurosci. 2017;11:9.

68. Bahar S, Suh M, Zhao M, Schwartz TH. Intrinsic optical signal imaging of neocortical seizures: the ’epileptic dip’. Neuroreport. 2006;17(5):499–503.

69. Zhao M, Ma H, Suh M, Schwartz TH. Spatiotemporal dynamics of perfusion and oximetry during ictal discharges in the rat neocortex. J Neurosci. 2009;29(9):2814–23.

70. Zhang C, Belanger S, Pouliot P, Lesage F. Measurement of Local Partial Pressure of Oxygen in the Brain Tissue under Normoxia and Epilepsy with Phosphorescence Lifetime Microscopy. PLoS One. 2015;10(8):e0135536.

71. Farrell JS, Gaxiola-Valdez I, Wolff MD, David LS, Dika HI, Geeraert BL, et al. Postictal behavioural impairments are due to a severe prolonged hypoperfusion/hypoxia event that is COX-2 dependent. Elife. 2016;5.

72. Colbourn R, Hrabe J, Nicholson C, Perkins M, Goodman JH, Hrabetova S. Rapid volume pulsation of the extracellular space coincides with epileptiform activity in mice and depends on the NBCe1 transporter. J Physiol. 2021;599(12):3195–220.

73. Fringuello AR, Colbourn R, Goodman JH, Michelson HB, Ling DSF, Hrabetova S. Rapid volume pulsations of the extracellular space accompany epileptiform activity in trauma-injured neocortex and depend on the sodium-bicarbonate cotransporter NBCe1. Epilepsy Res. 2024;201:107337.

74. Shulman RG, Rothman DL, Behar KL, Hyder F. Energetic basis of brain activity: implications for neuroimaging. Trends Neurosci. 2004;27(8):489–95.

75. Raichle ME, Mintun MA. Brain work and brain imaging. Annu Rev Neurosci. 2006;29:449–76.

76. Attwell D, Iadecola C. The neural basis of functional brain imaging signals. Trends Neurosci. 2002;25(12):621–5.

77. Hall CN, Howarth C, Kurth-Nelson Z, Mishra A. Interpreting BOLD: towards a dialogue between cognitive and cellular neuroscience. Philos Trans R Soc Lond B Biol Sci. 2016;371(1705).

78. Bolger FB, McHugh SB, Bennett R, Li J, Ishiwari K, Francois J, et al. Characterisation of carbon paste electrodes for real-time amperometric monitoring of brain tissue oxygen. J Neurosci Methods. 2011;195(2):135–42.

79. Dietzel I, Heinemann U, Hofmeier G, Lux HD. Transient changes in the size of the extracellular space in the sensorimotor cortex of cats in relation to stimulus-induced changes in potassium concentration. Exp Brain Res. 1980;40(4):432–9.

80. Dietzel I, Heinemann U, Lux HD. Relations between slow extracellular potential changes, glial potassium buffering, and electrolyte and cellular volume changes during neuronal hyperactivity in cat brain. Glia. 1989;2(1):25–44.

81. Sykova E, Nicholson C. Diffusion in brain extracellular space. Physiol Rev. 2008;88(4):1277–340.

82. de Lanerolle NC, Lee TS, Spencer DD. Astrocytes and epilepsy. Neurotherapeutics. 2010;7(4):424–38.

83. Murphy TR, Binder DK, Fiacco TA. Turning down the volume: Astrocyte volume change in the generation and termination of epileptic seizures. Neurobiol Dis. 2017;104:24–32.

84. Lauritzen M, Hansen AJ, Kronborg D, Wieloch T. Cortical Spreading Depression Is Associated with Arachidonic-Acid Accumulation and Preservation of Energy-Charge. J Cerebr Blood F Met. 1990;10(1):115–22.

85. Anwar F, Grech O, Mugo CW, Roberts JA, Hubbard JC, Thomas CN, et al. A systematic review of the causes and consequences of spreading depolarization in neuroinflammation; implications for neurovascular disorders. J Neuroinflammation. 2025;22(1):178.

86. Kramer DR, Fujii T, Ohiorhenuan I, Liu CY. Cortical spreading depolarization: Pathophysiology, implications, and future directions. J Clin Neurosci. 2016;24:22–7.

87. Kirchner T, Gröhl J, Herrera MA, Adler T, Hernández-Aguilera A, Santos E, Maier-Hein L. Photoacoustics can image spreading depolarization deep in gyrencephalic brain. Sci Rep-Uk. 2019;9.

88. von Bornstadt D, Houben T, Seidel JL, Zheng Y, Dilekoz E, Qin T, et al. Supply-demand mismatch transients in susceptible peri-infarct hot zones explain the origins of spreading injury depolarizations. Neuron. 2015;85(5):1117–31.

89. Somjen GG. Mechanisms of spreading depression and hypoxic spreading depression-like depolarization. Physiol Rev. 2001;81(3):1065–96.

90. Vorisek I, Sykova E. Ischemia-induced changes in the extracellular space diffusion parameters, K+, and pH in the developing rat cortex and corpus callosum. J Cereb Blood Flow Metab. 1997;17(2):191–203.

91. Windmuller O, Lindauer U, Foddis M, Einhaupl KM, Dirnagl U, Heinemann U, Dreier JP. Ion changes in spreading ischaemia induce rat middle cerebral artery constriction in the absence of NO. Brain. 2005;128(Pt 9):2042–51.

92. Ayata C, Lauritzen M. Spreading Depression, Spreading Depolarizations, and the Cerebral Vasculature. Physiol Rev. 2015;95(3):953–93.

93. Offenhauser N, Thomsen K, Caesar K, Lauritzen M. Activity-induced tissue oxygenation changes in rat cerebellar cortex: interplay of postsynaptic activation and blood flow. J Physiol. 2005;565(Pt 1):279–94.

94. Bickler PE, Buck LT. Adaptations of vertebrate neurons to hypoxia and anoxia: maintaining critical Ca2+ concentrations. J Exp Biol. 1998;201(Pt 8):1141–52.

95. Gnaiger E. Oxygen conformance of cellular respiration. A perspective of mitochondrial physiology. Adv Exp Med Biol. 2003;543:39–55.

96. Зыков С. RatSkull [Available from: https://stock.adobe.com/jp/images/old-rat-skull-close-up-on-black-background/276868267?prev_url=detail.

